# Primary souring: a novel bacteria-free method for sour beer production

**DOI:** 10.1101/121103

**Authors:** Kara Osburn, Justin Amaral, Sara R. Metcalf, David M. Nickens, Cody M. Rogers, Christopher Sausen, Robert Caputo, Justin Miller, Hongde Li, Jason M. Tennessen, Matthew L. Bochman

**Affiliations:** Molecular and Cellular Biochemistry Department, 212 South Hawthorne Drive, Simon Hall MSB1, room 405B, Indiana University, Bloomington, IN 47405, USA.; Mainiacal Brewing Company, Bangor, ME 04401, USA.; Wild Pitch Yeast, Bloomington, IN 47405, USA.; Department of Biology, Indiana University, 1001 East Third Street, Bloomington, IN 47405, USA.

**Keywords:** yeast, lactic acid, sour beer, heterolactic fermentation

## Abstract

In the beverage fermentation industry, especially at the craft or micro level, there is a movement to incorporate as many local ingredients as possible to both capture terroir and stimulate local economies. In the case of craft beer, this has traditionally only encompassed locally sourced barley, hops, and other agricultural adjuncts. The identification and use of novel yeasts in brewing lags behind. We sought to bridge this gap by bio-prospecting for wild yeasts, with a focus on the American Midwest. We isolated 284 different strains from 54 species of yeast and have begun to determine their fermentation characteristics. During this work, we found several isolates of five species that produce lactic acid and ethanol during wort fermentation: *Hanseniaspora vineae*, *Lachancea fermentati*, *Lachancea thermotolerans*, *Schizosaccharomyces japonicus*, and *Wickerhamomyces anomalus*. Tested representatives of these species yielded excellent attenuation, lactic acid production, and sensory characteristics, positioning them as viable alternatives to lactic acid bacteria (LAB) for the production of sour beers. Indeed, we suggest a new LAB-free paradigm for sour beer production that we term “primary souring” because the lactic acid production and resultant pH decrease occurs during primary fermentation, as opposed to kettle souring or souring via mixed culture fermentation.

**Chemical compounds studied in this article:** Lactic acid (PubChem CID: 612); Ethanol (PubChem CID: 702)

**Abbreviations:** ABV, alcohol by volume; DIC, differential interference contrast; EtOH, ethanol; FG, final gravity; gDNA, genomic DNA; IBU, international bittering unit; LAB, lactic acid bacteria; LASSO, lactic acid specific soft-agar overlay; N-J, neighbor-joining; OG, original gravity; WLN, Wallerstein Laboratories nutrient; YPD, yeast extract, peptone, and dextrose

## 1. Introduction

Currently, we are in the midst of a global craft beer boom, with the number of small independent breweries growing at a tremendous pace (1). This has led to increased competition, not only with the large macrobrewers but among the craft brewers themselves. As such, there is a need in the industry to differentiate oneself from, minimally, other local breweries. This has fueled experimentation with the core beer ingredients of water (2), malted grain (3), hops (4) and yeast (5), as well as with various adjuncts. Much of this experimentation is also focused on locally sourced ingredients to capture terroir and bolster the local economy (6,7).

Despite this widespread experimentation, the isolation and use of novel yeasts for brewing has lagged behind that of the other ingredients. This is in part due to the easy availability of numerous ale and lager strains from reputable commercial suppliers such as White Labs, Wyeast, and Lallemand (8). However, focusing on two species, *Saccharomyces cerevisiae* for ales and *Saccharomyces pastorianus* for lagers, naturally limits the genotypic and phenotypic variation available in brewing strains. This also translates into a limited palette of aromatic and flavor compounds made by these strains, especially considering their extremely high evolutionary relatedness (9,10).

To overcome this constraint, several laboratories and breweries have begun to culture wild yeasts and characterize their beer fermentation capabilities. Most efforts have focused on wild ale and lager strains (11,12) to increase the available genetic diversity of strains that naturally display high ethanol tolerance. However, multiple strains of yeasts in the *Brettanomyces*, *Hanseniaspora*, *Lachancea*, and *Pichia* genera (13-15) have also been investigated as alternative species for the production of beer.

We also recently began bio-prospecting for wild yeasts with desirable brewing characteristics (5). Here, we report the collection of nearly 300 strains from 26 genera. During trial wort fermentations, we found that strains from five species (*Hanseniaspora vineae*, *Lachancea fermentati*, *Lachancea thermotolerans*, *Schizosaccharomyces japonicus*, and *Wickerhamomyces anomalus*) were capable of heterolactic fermentation of sugar into lactic acid, ethanol, and CO_2_. Larger-scale brewing with four strains demonstrated that these yeasts are highly attenuative, flocculate well, yield appreciable levels of lactic acid, and produce pleasant aromatic and flavor compounds. We suggest a new paradigm for sour beer production called “primary souring” that avoids the use of lactic acid bacteria (LAB) and instead relies solely on lactic acid production by a heterofermentative yeast during primary fermentation.

## 2. Materials and methods

### 2.1. Strains, media, and other reagents

*S. cerevisiae* strain WLP001 was purchased from White Labs (San Diego, CA). Wild strains were isolated as described in (5). All yeast strains were routinely grown on yeast extract, peptone, and dextrose (YPD; 1% (w/v) yeast extract, 2% (w/v) peptone, and 2% (w/v) glucose) plates containing 2% (w/v) agar at 30°C and in YPD liquid culture at 30°C with aeration unless otherwise noted. All strains were stored as 15% (v/v) glycerol stocks at −80°C. Media components were from Fisher Scientific (Pittsburgh, PA, USA) and DOT Scientific (Burnton, MI, USA). All other reagents were of the highest grade commercially available.

### 2.2. Strain identification and phylogenetic analysis

To identify wild yeasts at the species level, frozen stocks were streaked onto YPD plates and incubated at 30°C until single colonies formed (18-48 h). Colonies were then picked into microcentrifuge tubes containing 100 μL of lysis solution (0.2 M LiOAc and 1% SDS) and incubated in a 65°C water bath for ≥15 min to lyse the cells. After 300 μL of 100% isopropanol was added to the tubes, they were mixed by vortexing, and the cell debris and genomic DNA (gDNA) were pelleted in a microcentrifuge for 5 min at maximum speed. The supernatant was decanted, and remaining traces were completely removed from the pellets by aspiration. The gDNA was resuspended in 50-100 μL TE buffer (10 mM Tris-HCl, pH 8, and 1 mM EDTA), and a 1-min spin at maximum speed was used to pellet the cell debris to clarify the DNA solution. The variable D1/D2 portion of the eukaryotic 26S rDNA was then amplified by PCR from the gDNA templates using oligos NL1 (GCATATCAATAAGCGGAGGAAAAG) and NL4 (GGTCCGTGTTTCAAGACGG) (11) and the following cycling conditions: 98°C for 5 min; 35 cycles of 98°C for 30 s, 55°C for 30 s, and 72°C for 30 s; and 72°C for 10 min. The PCRs were assessed for D1/D2 amplification by running 10% of the reaction volume on 1% (w/v) agarose gels at 100 V (560 bp expected product size). The amplified DNA was then purified using a PCR Purification Kit (Thermo Scientific, Waltham, MA) and quantified using a BioTek Synergy H1 plate reader. The DNA was sequenced by ACGT, Inc. (Wheeling, IL) using primer NL1, and the sequence was used to query the National Center for Biotechnology Information nucleotide database with the Basic Local Alignment Search Tool (BLAST; http://blast.ncbi.nlm.nih.gov/Blast.cgi?CMD=Web&PAGE_TYPE=BlastHome).

After species identification, the phylogenetic relationships among the isolated strains of *H. vineae*, *L. fermentati*, *L. thermotolerans*, *S. japonicus*, and *W. anomalus* were determined by aligning their 26S rDNA sequences using ClustalX (16). The alignments were iterated at each step but otherwise utilized default parameters. ClustalX was also used to draw and bootstrap neighbor-joining (N-J) phylogenetic trees using 1000 bootstrap trials; the trees were visualized with TreeView v. 1.6.6 software (http://taxonomy.zoology.gla.ac.uk/rod/rod.html). The *Schizosaccharomyces pombe* rDNA sequence (GenBank accession HE964968) was included in the alignments as the outgroup, and this was used to root the N-J tree in TreeView. WLP001 was included to determine the relatedness of the wild strains to a commercially available ale yeast.

### 2.3. Test fermentations

For laboratory-scale fermentations, select yeast strains were streaked for single colonies onto YPD plates as described above and grown to saturation in 4 mL of YPD liquid medium overnight at 30°C with aeration. The cell count of the starter cultures was approximated by measuring the OD_660_ and converting that value to cells/mL as described at http://www.pangloss.com/seidel/Protocols/ODvsCells.html. In most cases, the saturated overnight cultures reached densities of ~5×10^8^ cells/mL. These starter cultures were then used to inoculate ~400 mL of blonde ale wort in 500 mL glass bottles capped with drilled rubber stoppers fitted with standard plastic airlocks. The wort was produced by mashing 65.9% Pilsner (2 Row) Bel and 26.9% white wheat malt at 65°C (149°F) for 75 min in the presence of 1 g/bbl CaCO_3_ and 1.67 g/bbl CaSO_4_ to yield an original gravity (OG) of 1.044. During the boil, 7.2% glucose was added, as well as Saaz hops to 25 international bittering units (IBUs). The fermentation cultures were incubated at 22.3±0.3°C (~72°F) for 2 weeks. Un-inoculated wort was treated as above to control for wort sterility. Prior to bottling into standard 12-oz brown glass bottles, their final gravity (FG) was measured using a MISCO digital refractometer (Solon, OH), and pH was measured using an Accumet AB150 pH meter (Fisher Scientific). Bottle conditioning was conducted as in (17) at room temperature for ≥2 weeks.

Small-batch fermentations were performed at Mainiacal Brewing in Bangor, ME. To produce the test wort, 93.4% two-row base malt and 6.6% carapils were mashed at 66.7°C (152°F) to yield an OG of 1.046. During the boil, Loral hops were added to a final concentration of 5.3 IBUs. The wort was then chilled and split into 5-gal portions in separate carboys. Approximately 1x10^11^ cells of the indicated yeast strains were used to inoculate the carboys and allowed to ferment under anaerobic conditions at 21.7°C (71°F) for 1 month. Gravity measurements were taken both with a hydrometer and refractometer by standard methods. The final pH was recorded prior to bottling and bottle conditioning as above.

### 2.4. Lactic acid specific soft-agar overlay (LASSO)

The production of lactic acid by yeast cells was assayed as described in (18). Briefly, cells were grown overnight in liquid YPD medium at 30°C with aeration. Then, 2 μL of each culture was spotted onto YPD10 plates (YPD agar supplemented with glucose to a final concentration of 10% w/v), allowed to absorb, and incubated overnight at 30°C. The plates were then covered with 6.5 mL soft-agar (0.5% agar in 300 mM Tris and 187 mM glutamate, pH 8.3). Upon solidification of the soft agar, a second soft-agar overlay was prepared by mixing 3.2 mL 1% agar with 3.2 mL of a staining solution composed of 30 mM Tris/18.75 mM glutamate (pH 8.3), 2.5 mM NAD, 0.5 mg/mL nitrotetrazolium blue, 125 μg/mL phenazine methosulfate, 7 U glutamate pyruvate transaminase, and 7 U L[+]-lactate dehydrogenase (all LASSO components were from Sigma-Aldrich, St. Louis, MO). Strains producing lactic acid formed purple halos within 10 min, the color of which darkened with increasing incubation time at room temperature. Cells not producing lactic acid never formed halos.

### 2.5. Multi-well lactic acid production assay

Because only a small number of strains can be tested for lactic acid production on a single plate in the LASSO assay, we also adapted it for use in multi-well plates. Briefly, individual strains were grown overnight in 100 μL YPD10 medium in the wells of 96-well plates at 30°C with aeration in a BioTek Synergy H1 plate reader. To avoid evaporation of the medium, 50 μL mineral oil was used to overlay each well. Then, 100 μL of staining solution (30 mM Tris/18.75 mM glutamate (pH 8.3), 2.5 mM NAD, 0.5 mg/mL nitrotetrazolium blue, 125 μg/mL phenazine methosulfate, 1 U/mL glutamate pyruvate transaminase, and 1 U/mL L[+]-lactate dehydrogenase) was added to each well and mixed by agitation in the plate reader. The reaction proceeded for ≥10 min at room temperature, and the presence of lactic acid was indicated by the gold colored solution turning green (and eventually blue with extended incubation).

### 2.6. Beer sensory analysis

Sensory analysis was performed by 10 volunteers with various levels of experience, from neophytes to those with Beer Judge Certification Program (19) training. In all cases, the sensory descriptors (*e.g*., Barnyard, Bitter, Body, Drinkability, Dry, Estery, Harsh, Hoppy, Malty, Papery, Sour, Sulfury, and Sweet in Supplemental Fig. S3) were defined and described to the participants, and commercial calibrations beers were used as examples of sour (Cauldron; Upland Brewing Company, Bloomington, IN, USA) and clean beers (Dragonfly IPA; Upland Brewing Company). Then, chilled samples of the experimental beers were provided to the volunteers, and they were instructed to write down aroma and flavor descriptions, as well as to rank each of the Supplemental Figure S3 flavor descriptors on a 1-10 scale. These analyses were performed in a blinded manner, with none of the participants knowing which yeast strains fermented the beers. After individual assessments, the group discussed the sensory attributes of each beer to come to a consensus on the best descriptions of aroma and flavor, which are reported throughout this work.

### 2.7. Gas chromatography-mass spectrometry (GC-MS) analysis of lactic acid

To determine the concentration of lactic acid in beer samples, 20 μL of beer and lactate standards were individually added to 900 μL of 90% methanol containing 1.25 μg/mL succinic-d4 acid in 1.5-mL microfuge tubes. The samples were vortexed for 10 s, incubated for 1 h at - 20ºC, and centrifuged at 20,000 x g for 5 min at 4ºC. The cleared supernatants were transferred to 1.5-mL tubes and dried overnight using a vacuum centrifuge (Savant). Subsequent derivatization and GC-MS analysis were performed as previously described (20). Lactate concentration was determined using a standard curve plotted from the analyzed lactate standards of known concentration.

## 3. Results

### 3.1. Discovery of five heterofermentative yeast species

We previously reported an initial description of our bio-prospecting and characterization of 100 wild yeasts for use in the brewing industry (5). We have increased our collection to 284 strains from 54 different species in 26 genera (Supplemental Table 1). To determine the relative usefulness of these strains in beer brewing, small laboratory-scale beer fermentations were performed for each isolate. During our sensory analyses of the resultant beers, we noted that many of the strains were producing beers that were characterized as tart or sour (Table 2), akin to styles that are produced with the aid of LAB (21). Initially, we hypothesized that these experimental beers may have been contaminated by LAB. However, when we phylogenetically grouped these strains, we found that they were all members of five species in four genera: *H. vineae*, *L. fermentati*, *L. thermotolerans*, *S. japonicus*, and *W. anomalus* (Supplemental Table 1 and Fig. 1). Because these strains were not randomly distributed throughout the many species in our collection, we hypothesized that the yeasts may be producing the lactic acid themselves. To determine if this apparent heterofermentative activity was specific to evolutionarily closely related yeasts, we aligned the sequences of the D1/D2 variable region of their ribosomal DNA and plotted a phylogenetic tree. As shown in Figure 1 and Figure S1, three of the species (*H. vineae*, *L. fermentati*, and *L. thermotolerans*) are closely related to ale yeast (WLP001), but the other two (*S. japonicus*, and *W. anomalus*) form more distinct clades, with the divergence between budding yeasts such as *S. cerevisiae* (*e.g*., WLP001) and fission yeasts such as *S. japonicus* (*e.g*., YH156) occurring approximately 1 billion years ago (22).

**Table 1.** Wild yeast species diversity and number of isolates.

| **Genus** | **Species** | **Isolates** |
| --- | --- | --- |
| *Aureobasidium* | *pullulans* | 2 |
| *Brettanomyces* | *bruxellensis* | 7 |
| *Candida* | *carphophila* | 1 |
|  | *intermedia* | 1 |
|  | *sake* | 2 |
|  | *tropicalis* | 6 |
|  | *zemplanina* | 1 |
|  | *zeylandoides* | 1 |
| *Clavispora* | *lusitaniae* | 3 |
| *Cryptococcus* | *albidus* | 1 |
| *Cyberlindnera* | *fabianii* | 5 |
|  | *rhodanensis* | 1 |
| *Debaryomyces* | *hansenii* | 7 |
| *Hanseniaspora* | *opuntiae* | 1 |
|  | *uvarum* | 12 |
|  | *valbyensis* | 1 |
|  | *vineae* | 11 |
| *Issatchenkia* | *orientalis* | 1 |
|  | *terricola* | 3 |
| *Kazachstania* | *unispora* | 1 |
| *Kluyveromyces* | *lactis* | 10 |
|  | *marxianus* | 10 |
| *Kodamaea* | *ohmeri* | 2 |
| *Kwoniella* | *mangroviensis* | 1 |
| *Lachancea* | *fermentati* | 8 |
|  | *kluyveri* | 4 |
|  | *thermotolerans* | 25 |
| *Metchnikowia* | *fruiticola* | 1 |
|  | *pulcherrima* | 1 |
| *Meyerozyma* | *guilliermondii* | 4 |
| *Ogatea* | *naganishii* | 1 |
| *Pichia* | *fermentans* | 3 |
|  | *galeiformis* | 12 |
|  | *guilliermondii* | 2 |
|  | *kluyveri* | 7 |
|  | *kudriavzevii* | 9 |
|  | *manshurica* | 4 |
|  | *membranifaciens* | 5 |
|  | *mexicana* | 2 |
|  | *nakazawae* | 2 |
|  | *quercuum* | 1 |
| *Rhodosporidium* | *babjevae* | 1 |
|  | *diobovatum* | 1 |
| *Rhodotorula* | *mucilaginosa* | 1 |
| *Saccharomyces* | *cerevisiae* | 37 |
|  | *kudriavzevii* | 1 |
|  | *paradoxus* | 8 |
| *Schizosaccharomyces* | *japonicus* | 2 |
|  | *pombe* | 4 |
| *Starmerella* | *bacillaris* | 2 |
|  | *bombicola* | 3 |
| *Torulaspora* | *delbrueckii* | 24 |
| *Wickerhamomyces* | *anomalus* | 17 |
| *Williopsis* | *saturnus* | 1 |

**Table 2.** Lab-scale fermentation and sensory results for representative heterofermentative strains. The highlighted strains were used in Figure 2.

| Strain | Species<sup>a</sup> | Place of origin | Collected from | Attenuation<sup>b</sup> | Final pH<sup>c</sup> | Sensory notes |
| --- | --- | --- | --- | --- | --- | --- |
| WYP39 | *L. fermentati* | Evansville, IN | Birch bark | 60% | 3.68 | Sour, pear, melon, black tea |
| YH25 | *L. fermentati* | New River Gorge, WV | Red oak bark | 60% | 3.66 | Lactic tart finish |
| YH26 | *L. thermotolerans* | New River Gorge, WV | Tulip poplar bark | 60% | 3.35 | Very tart, peach, citrus zest |
| YH27 | *L. thermotolerans* | New River Gorge, WV | Red oak bark | 40% | 3.42 | Tart green apple, clean |
| YH72 | *H. vineae* | New Kensington, PA | Ash bark | 75% | 3.26 | Slightly sour, clean, tart fruit, apple cider, quaffable |
| YH73 | *L. thermotolerans* | New Kensington, PA | Mulberries | 55% | 3.36 | Very sour, berries |
| YH77 | *L. fermentati* | Princeton, NJ | Red oak bark | 60% | 3.55 | Tart, clean, pear |
| YH79 | *L. thermotolerans* | Holmdel, NJ | Red oak bark | 50% | 3.42 | Slightly tart, clean, rounded |
| YH81 | *L. thermotolerans* | Holmdel, NJ | White oak bark | 55% | 3.21 | Tart, fruity, citrus, pear, blood orange |
| YH82 | *W. anomalus* | Sarver, PA | Shumard oak bark | 83% | 3.24 | Slightly sour, fruity, clean, aromatic, tart fruit, reminiscent of perry |
| YH109 | *L. thermotolerans* | Indianapolis, IN | Bell pepper | 55% | 3.39 | Very sour, clean |
| YH140 | *L. thermotolerans* | Bloomington, IN | Shagbark hickory bark | 50% | 3.26 | Sour, clean, balanced, tea flavors |
| YH156 | *S. japonicus* | Northeastern PA | Oak bark | 72% | 3.88 | Sour, fruity, green apple Jolly Rancher |

<sup>a</sup> The genus abbreviations are: *L.*, *Lachancea*; *H.*, *Hanseniaspora*; *W.*, *Wickerhamomyces*; and *S.*, *Schizosaccharomyces*.
<sup>b</sup> The reported attenuation is based on several trial fermentations in various worts.
<sup>c</sup> The initial pH was 5.0.

**Figure 1.**
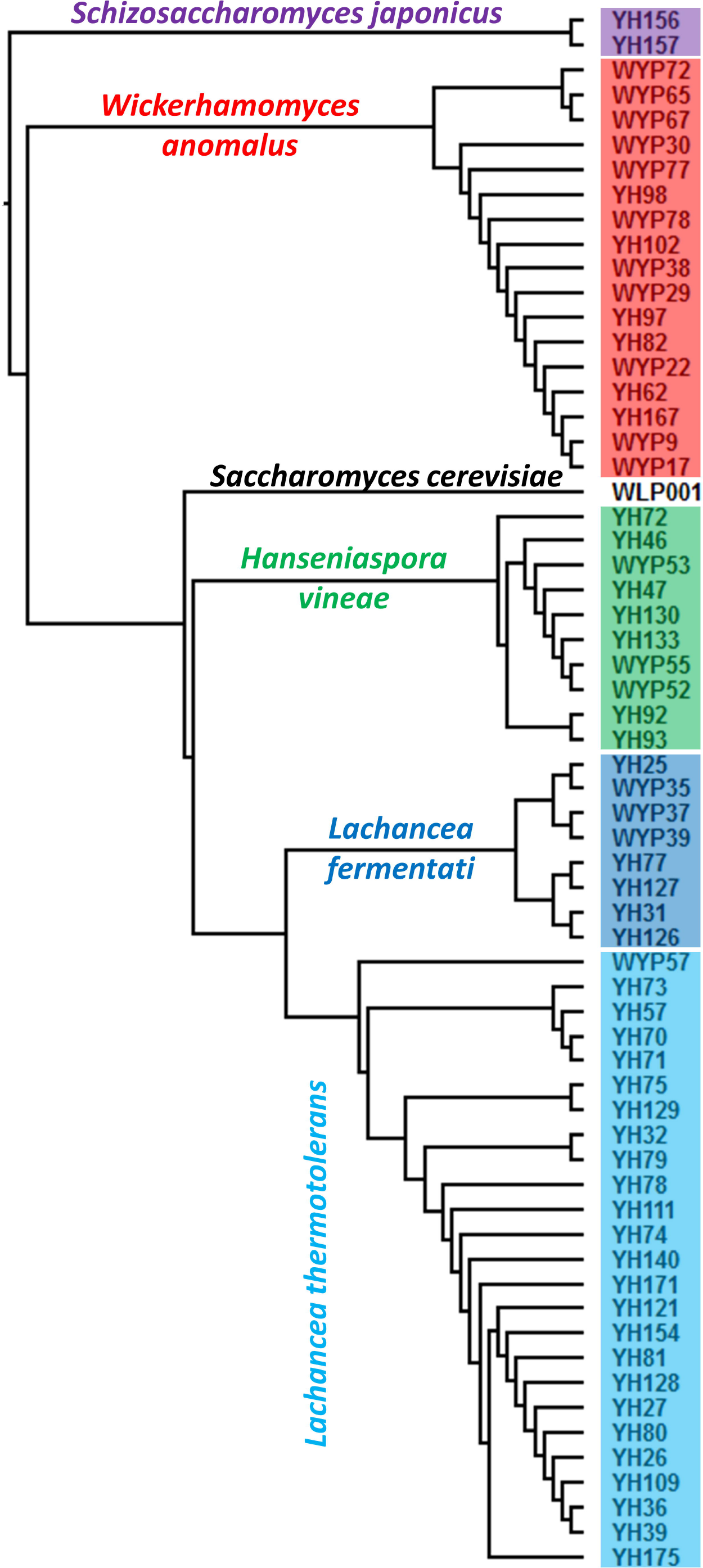
Evolutionary relationships among the wild strains and a commercially available ale yeast. The D1/D2 rDNA sequences of the indicated strains were aligned, and the phylogenetic relationships among them were drawn as a rooted N-J tree using *Schizosaccharomyces pombe* as the outgroup. From top to bottom, the *S. japonicus* strains are highlighted purple, the *W. anomalus* strains are red, the *H. vineae* strains are green, the *L. fermentati* strains are dark blue, and the *L. thermotolerans* strains are light blue. *S. cerevisiae* strain WLP001 is not highlighted and occupies the relative midpoint of the phylogenetic tree.

Regardless of their evolutionary relationships, the strains listed in Table 2 and other isolates of the same species (data not shown) varied in their fermentative activities. Their levels of attenuation varied from 40-83%, with decreases in the initial pH of 5.0 to as low as 3.21. Although some of these differences may be attributable to differences among the species themselves, intra-species differences were also noted, especially during sensory analyses. For instance, *L. thermotolerans* YH73 produced a “very sour” flavor with berry notes, but the same beer fermented with *L. thermotolerans* YH79 was characterized as only “slightly tart” yet clean and rounded (Table 2).

*3.4. Strains* L. fermentati *WYP39*, H. vineae *YH72*, W. anomalus *YH82*, L. thermotolerans *YH140, and* S. japonicus YH156 *produce lactic acid*

We further sought to determine if strains *L. fermentati* WYP39, *H. vineae* YH72, *W. anomalus* YH82, *L. thermotolerans* YH140, and *S. japonicus* YH156 were truly producing lactic acid during fermentation, rather than one or more other secondary metabolites that yield a tart/sour flavor (23). Using the LASSO assay for lactic acid production by yeast (18), we found that all five strains produced lactic acid (denoted by dark halos in Fig. 2A), similar to the *Lactobacillus plantarum* positive control. In contrast, the *S. cerevisiae* WLP001 negative control failed to generate a halo. Because the LASSO assay can only be used for a limited number of strains on a single plate, we adapted it into a multi-well plate assay (Fig. 2B). Here, lactic acid production is evident by the golden-colored assay medium turning green, as indicated by the LAB controls. Again, WLP001 failed to generate detectable lactic acid, as did a common research strain of *E. coli*. However, multiple tested isolates of *L. thermotolerans*, *L. fermentati*, *H. vineae*, *S. japonicus*, and *W. anomalus* did test positive for lactic acid production. Some individual *L. thermotolerans* strains failed to generate lactic acid (wells 2 and 4) or did so slowly, as certain wells were just beginning to turn green (well 3) when the image in Figure 2B was acquired. These results correspond with sensory analysis of beers fermented with the various *L. thermotolerans* strains, which ranged from not sour to very tart (Table 2 and data not shown).

**Figure 2.**
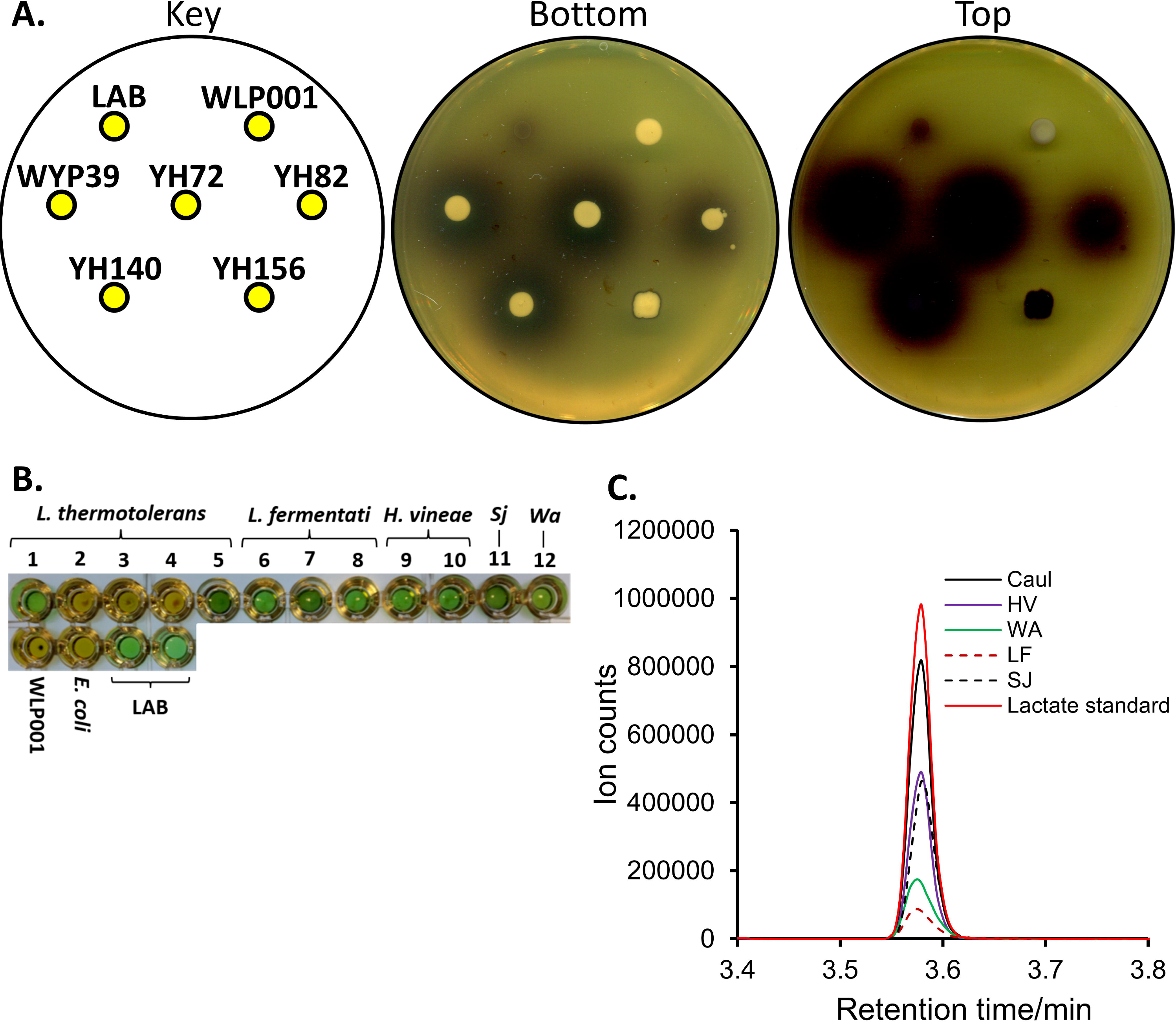
Lactic acid production. A) LASSO assay for lactic acid production by the indicated strains. Cells producing lactic acid develop a dark halo. Images of both the top and bottom of a representative LASSO plate are shown. *S. cerevisiae* WLP001 and LAB (*L. plantarum*) were included as negative and positive controls for lactic acid production, respectively. These results are indicative of three independent experiments using the same strains. B) Multi-well plate assay for lactic acid production. Cell growth medium containing lactic acid turns from gold to green when the enzymatic assay is complete. Multiple strains of *L. thermotolerans, L. fermentati, H. vineae*, and LAB were tested, as well as *S. japonicus* YH156 (*Sj*), *W. anomalus* YH82 (*Wa*), *S. cerevisiae* WLP001, and *Escherichia coli* DH5α (*E. coli*). WLP001 and *E. coli* served as negative controls for lactic acid production by yeast and bacteria, respectively. *L. plantarum* (left LAB well) and *L. brevis* (right LAB well) were used as positive controls for lactic acid production. C) L-lactate quantification. Typical GC-MS spectra of the lactate standard (solid red line) and isolated lactate from beer samples are shown. HV (solid purple line), WA (solid green line), LF (dashed burnt sienna line), and SJ (dashed black line) represent beers fermented with *H. vineae* YH72, *W. anomalus* YH82, *L. fermentati* WYP39, and *S. japonicus* YH156, respectively. “Caul” (solid black line) represents the lactate isolated from Cauldron, a positive control for lactate in beer (See Section 2.6 and (17)).

### 3.4. Analysis of beers fermented with lactic acid-producing yeasts

To monitor the activities of *L. fermentati* WYP39, *H. vineae* YH72, *W. anomalus* YH82, and *S. japonicus* YH156 in larger-scale fermentations, we inoculated these strains into glass carboys containing 19 L (5 gal) each of an identical blonde wort. Because the beer brewing capabilities of a *L. thermotolerans* strain were recently described (14), we omitted strain *L. thermotolerans* YH140 from these assays to avoid generating redundant data. All of the strains had short lag times (*i.e*., the approximate time from inoculation to the first visible signs of fermentation) ranging from 6-14 h (Table 3). These lag times to fermentation were similar to that of WLP001 inoculated in a similar blonde wort (~12 h, Table 3). However, the kinetics of the full fermentation differed for each lactic acid-producing yeast. *L. fermentati* WYP39 had the shortest lag time and fermented rapidly for 2 weeks, at which point it slowed considerably and required an additional 2 weeks to reach a final gravity of 0.099 (Table 3). *H. vineae* YH72 was a slow and steady fermenter, attaining a terminal gravity of 1.000 after 3 weeks. *W. anomalus* YH82 was similar, but required a full 4 weeks to reach a final gravity of 1.001.

**Table 3.** Large-scale fermentation data for select heterofermentative yeasts and the WLP001 control.

| Strain | Lag time | FG<sup>b</sup> | Final pH<sup>c</sup> | Flocculation | Sensory |
| --- | --- | --- | --- | --- | --- |
| WYP39<sup>a</sup> | 6 h | 0.099 | 3.74 | Medium | Tart, dry, light pineapple & mango |
| YH72 | 14 h | 1.000 | 3.23 | Medium | Very sour, stone fruit flavors |
| YH82 | 13 h | 1.001 | 3.36 | Medium | Very sour, pear, apple, and apricot |
| YH156 | 14 h | 0.099 | 3.20 | High | Sour, intense stone fruit aroma & flavors |
| WLP001 | 12 h | 1.010 | 4.35 | Medium | Neutral, clean, slightly fruity |

<sup>a</sup> WYP39 = *Lachancea fermentati*, YH72 = *Hanseniaspora vineae*, YH82 = *Wickerhamomyces* *anomalus*, YH156 = *Schizosaccharomyces japonicus*, and WLP001 = *Saccharomyces cerevisiae*.
<sup>b</sup> FG, final gravity; original gravity = 1.046 for all fermentations.
<sup>c</sup> The starting pH was 5.35 for all fermentations.

The *S. japonicus* YH156 strain displayed the most variant fermentative characteristics. After reaching a vigorous state of fermentation at 14 h (Table 3), popcorn-like clusters of cells formed and floated around within the fermenter (Supplemental Fig. 2). Eventually, they settled into a mountainous pile against one side of the carboy before compacting down into a yeast slurry with a typical appearance on the bottom of the fermenter. The fermentation reached a final gravity of 0.099 approximately 27 days after inoculation. The final pH of each beer was recorded and varied from a low of 3.20 to a high of 3.74. Sensory analyses of each beer were conducted (Supplemental Fig. 3), and the tasting notes are discussed in Sections 4.1-4.5 below.

We also quantified the concentration of L-lactic acid present in each beer by GC-MS (Fig. 2C and S4). We used Cauldron, a commercially available sour beer made by mixed fermentation of yeast and LAB (17), as a positive control for lactic acid production; it contained 100.54 mM lactate. Based on a lactate standard curve, *L. fermentati* WYP39, *H. vineae* YH72, *W. anomalus* YH82, and *S. japonicus* YH156 produced 10.02, 35.69, 29.05, and 50.09 mM lactate, respectively.

The *S. japonicus* YH156 results contrast with those from the LASSO assay in Figure 2A, where *S. japonicus* YH156 displayed the least evidence of lactic acid production. Because the lactic acid production occurred in the presence of oxygen in the LASSO assay, this may indicate that *S. japonicus* YH156 is Crabtree negative or only weakly Crabtree positive, *i.e*., requiring anaerobic conditions for fermentation (see (24) and references therein). It should also be noted that the beers analyzed by GC-MS were fermented in a brewery that utilizes LAB, and thus, it remains a formal possibility that they may have been inadvertently contaminated by other organisms that can generate lactic acid. However, the results in Figures 2A and 2B are from pure cultures (as judged by post-fermentation plating, microscopy, and PCR analyses of the yeast slurry for LAB contamination; Supplemental Fig. 5 and data not shown) of *L. fermentati* WYP39, *H. vineae* YH72, *W. anomalus* YH82, and *S. japonicus* YH156. These cultures also still acidified beer during fermentation in the presence of antibiotics or 75 IBU wort in a laboratory setting (Supplemental Table 2), strongly suggesting that LAB contamination was not the source of the lactic acid.

## 4. Discussion

The next phase in the “local” movement in the beer industry will be the isolation and use of local yeast strains in brewing. Indeed, in the U.S., nearly all commercially available ale and lager strains are of European origin, so no American beer will ever truly be local without the inclusion of New World yeast. Here, we detailed the initial characterization of nearly 300 local strains for use in fermentation. Within this strain bank, we uncovered five species that generate lactic acid and ethanol during primary fermentation and suggest that they can be used in a novel, LAB-free beer souring method (see Section 4.6. below).

### 4.1. H. vineae

To our knowledge, this is the first report of pure cultures of *H. vineae* being used to ferment beer. As the species name suggests, *H. vineae* is typically associated with wine, where it has previously been investigated alone and in combination with *S. cerevisiae* for grape must fermentation (25-27). The strains previously tested are notable for the production of high levels of 2-phenylethyl acetate, which is an aromatic compound that lends floral, fruity, and/or honey-like notes to wine (26). Indeed, in our trial fermentations with *H. vineae* YH72, we often noted fruity aromas and flavors (Tables 2 and 3).

Although yeasts in the *Hansenia* genus are the predominant species found on grapes (28-30), they are also found elsewhere (reviewed in (31)). Strain *H. vineae* YH72 was isolated from a white ash tree (*Fraxinus americana*) in southwestern Pennsylvania (Table 2) and was the only *Hanseniaspora* isolate to consistently produce tart beer (see Supplemental Table 3). It ferments slowly relative to typical commercially available ale yeasts, but reached high levels of attenuation after only two weeks (Table 2) and further attenuated with additional fermentation time (Table 3). The beers produced by short fermentations with *H. vineae* YH72 were slightly sour but clean and highly drinkable, with notes of apple cider. Longer fermentation yielded very sour beer with a pH (3.23) and acidic bite reminiscent of beers produced with LAB, as well as stone fruit notes (Table 3, Supplemental Fig. 3). Currently, we are further characterizing the fermentative capabilities and acid production of *H. vineae* YH72 and other *Hansenia* species.

**Figure 3.**
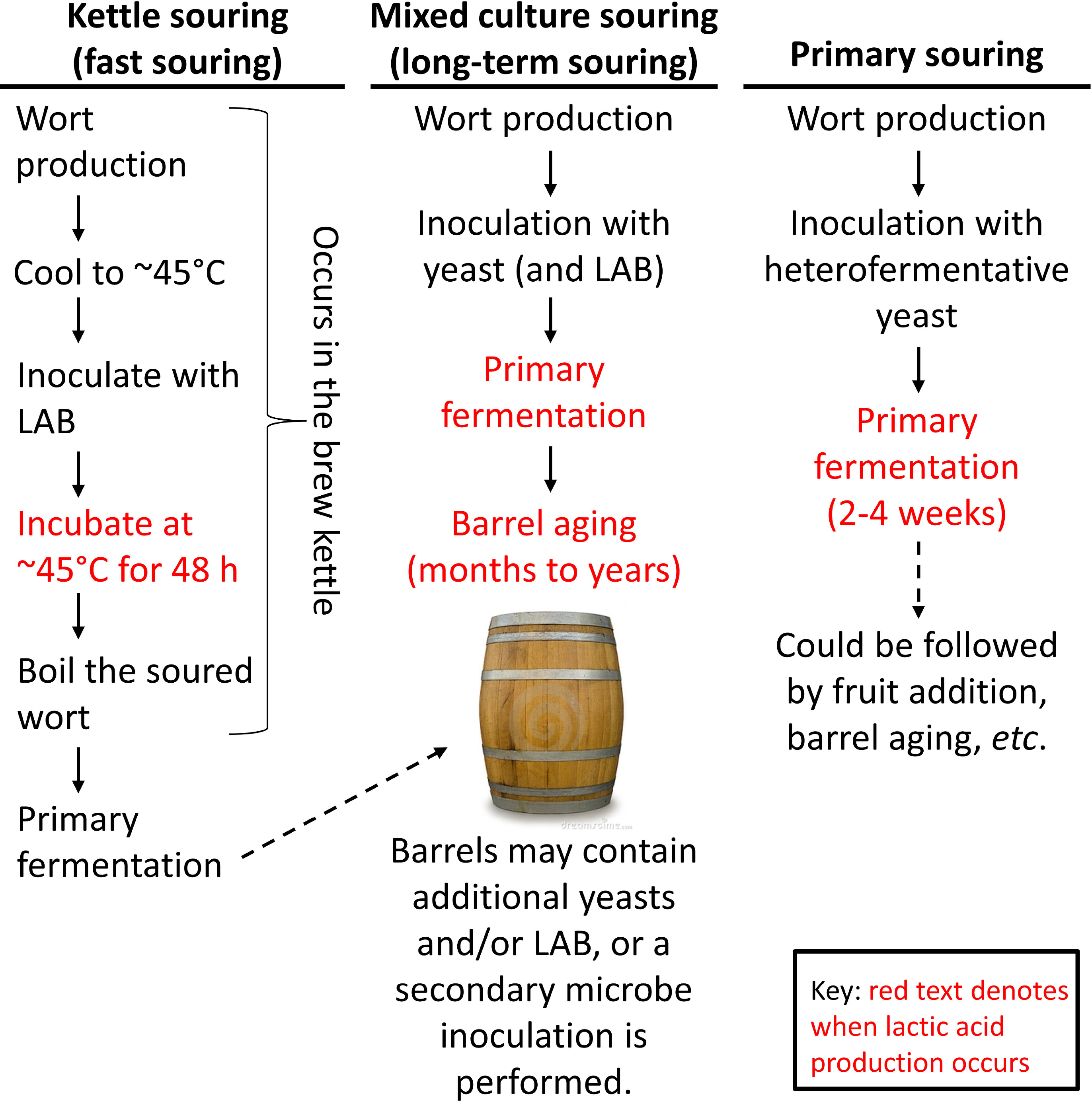
Comparison of kettle souring, wood-aged souring, and primary souring. Left) During kettle souring, unhopped or lightly hopped wort is produced in a brew kettle as normal, but then it is only partially cooled to approximately 40-45°C. This temperature favors the growth of LAB, which can be introduced by inoculation with pure cultures or the addition of grain. The LAB then sour the wort in the brew kettle to the desired pH, which usually occurs over the course of 24-48 h. The soured wort in the kettle is ultimately boiled a second time to kill the LAB, and hops can be introduced at this point. The wort is then transferred to a fermenter and (typically) inoculated with *S. cerevisiae* for primary fermentation. Middle) During mixed culture souring, lightly hopped wort is produced in the brew kettle and transferred to a fermenter. There, it can be inoculated with *S. cerevisiae* for primary fermentation. In some cases, LAB are added prior to or concurrently with *S. cerevisiae* (or *Brettanomyces* spp. in 100% “Brett” beers). If the LAB are added at this stage, souring begins during primary fermentation. After the yeast has attenuated the beer to the desired level, it is then barrel aged for months or years until it attains a low pH and complex flavor profile. Barrel aging is another stage at which LAB and *Brettanomyces* spp. can be added (either resident in the barrels or as pure inocula) to induce souring. Right) Primary souring: see the text in Section 4.6 for details.

### 4.2. L. fermentati

Very little is known about *L. fermentati*, especially with regard to beverage fermentation. Industrially, this species of yeast has been found in fermented (wine (32), cachaça (33), and water kefir (34)) and non-fermented beverages (coconut water and fruit juices (35)). However, its effects on the sensory characteristics of these beverages are largely unknown. As with *H. vineae* YH72 above, this also appears to be the first report of beer fermented with pure cultures of *L. fermentati*. Our laboratory-scale test fermentations indicated that strain WYP39 displayed decent wort attenuation for a wild strain (60%, Table 2), and longer fermentation in a larger-scale fermentation yielded a dry product (Table 3). The final pH was modest compared to other sour beers (Table 3), creating a flavor that was more tart than sour, but this was accentuated by light pineapple and mango flavors. There are many species in the *Lachancea* genus (36). Based on the desirable brewing characteristics of *L. fermentati* and *L. thermotolerans* (discussed below in Section 4.3. and in Domizio *et al*. (14)), it will be interesting to assess the activities of these other species during beer fermentation.

### 4.3. L. thermotolerans

Various strains of *L. theromtolerans* have been studied for their effects on wine fermentation (reviewed in (29)), though usually in co-fermentations with *S. cerevisiae* (*e.g*., (37,38). Recently, *L. thermotolerans* strain Lt101 was shown to be a viable yeast for beer production (14), and the same group also found that three *L. thermotolerans* strains including Lt101 produce lactic acid during fermentation. Although this is similar to our observations (Table 2 and data not shown), the strains investigated by Domizio *et al*. (14) only decreased the pH of wort from 5.66 to 4.28-3.77 during fermentation. Most of their experiments yielded final pH values in the 4.17-4.3 range, however, which is similar to the pH decrease caused by *S. cerevisiae* UCD 915. Our *L. thermotolerans* isolates that produced noticeably tart beers reached terminal pH values of ~3.35 (Table 2 and data not shown). These discrepancies are likely due to differences in the experimental set ups, as well as strain-to-strain variability, which is discussed in Section 5 below.

### 4.4. S. japonicus

*S. japonicus* is closely related to *S. pombe*, which was originally isolated from African millet beer (reviewed in (39)). *S. japonicus* itself was first isolated from strawberries in Japan (40) and is associated with indigenous fermented beverages (*e.g*., kaffir beer, plantain beer, palm wine, sugar cane wine, and sake) around the world (41), wine production (42), and was isolated from spontaneously fermented beer in North Carolina (43). However, no characterization of that beer is available. Thus, this is the first rigorous report of *S. japonicus* used for primary fermentation of beer. The attenuation levels of strain *S. japonicus* YH156 were excellent in both laboratory- and large-scale fermentations (Table 2 and 3), and the aroma and flavor profiles of these beers included common descriptors of sour, fruity, and stone fruit. Individual tasters identified green apple Jolly Rancher, tart apple, pear, pineapple, and peach notes.

### 4.5. W. anomalus

*W. anomalus*, previously known as *Saccharomyces anomalus* (44), has multiple roles in the biotechnology, agriculture, and food production fields. It is often found in association with grain and is especially useful to inhibit storage molds during malting (45). Concerning beverage fermentation, *W. anomalus* is generally referred to as a beer spoilage organism (46), but it is also necessary for cocoa and coffee bean fermentation (47). Although *W. anomalus* has been investigated for its use in apple wine and hard cider production (48,49), the only work involving beer has focused on the spoilage properties of this species (46). As with *S. japonicus* YH156 above, the YH82 strain of *W. anomalus* yielded excellent attenuation at both fermentation scales tested (Tables 2 and 3). This strain produced a less intense sour character than others, but the beer was characterized as clean, aromatic, and fruity with notes of pear, apple, and apricot.

### 4.6. Primary souring

There are two general methods by which sour beers are produced: kettle souring and mixed culture fermentation (Fig. 3) (21). Kettle souring is the more rapid and modern technique. This method affords brewers tight control over acid production (souring can be stopped at any time via boiling) and is less time consuming that mixed culture fermentation (below), but it also has inherent weaknesses. First, the entire souring process occurs in the brew kettle, so it prevents additional wort production in that vessel. Indeed, kettle souring is often relegated to weekends when small breweries are otherwise not in production mode. Second, boiling the wort after it has been soured drives off volatile aromatics that may also have been produced during souring, yielding beers that are described and criticized as lacking in depth and character. To combat some of these sensory downsides, some brewers are now barrel aging kettle sours to impart oak complexity to the final beers.

In contrast, souring by mixed culture fermentation is the more traditional method. It produces more complex and nuanced flavors in beer than kettle souring, but it suffers from a huge time lag from wort production to the final beer and requires a large space dedicated to housing the barrels. Further, the souring organisms are usually alive throughout the beer production and packaging processes, so separate equipment is usually necessary to prevent the accidental contamination of non-sour (*i.e*., clean) beers.

Here, we propose a third paradigm for sour beer production that we call primary souring. In this method, the wort is inoculated with a yeast capable of heterolactic fermentation rather than *S. cerevisiae*, and souring occurs during primary fermentation in the absence of LAB (Fig. 3). The yeast strains described in Tables 2 and 3 and in the text above did not display rapid fermentation kinetics like commercially available ale yeasts, but they still completed fermentation within a month, displaying excellent levels of attenuation and medium-to-high flocculation (Table 3). Further, the sensory profiles of the beers were superior to kettle soured beer, displaying both lactic tartness and fruity aromatic and flavor notes. Compared to the sour production methods above, primary souring is beneficial in that it frees up the brew kettle and does not require lengthy aging in barrels, though oak aging is a possibility after primary fermentation (Fig. 3). Perhaps most alluringly, primary souring does not require the introduction of bacteria into the brewery, and preliminary tests suggest that *H. vineae*, *L. fermentati*, *L. thermotolerans*, *S. japonicus*, and *W. anomalus* can be eliminated as easily as *S. cerevisiae* from brewing equipment using standard clean-in-place protocols. As is typical of yeasts, these species are also hop tolerant (Supplemental Table 2), enabling more liberal use of hops in wort production for sour beers. However, it should be noted that yeast growth can be inhibited by hop iso-α-acids in acidic milieus (50), so the absolute levels of hop tolerance will likely vary by strain and the desired pH of the sour beer.

## 5. Conclusions and Outlook

We set out to isolate and characterize new yeasts for use in beer fermentation. Here, we highlight the discovery of a set of yeast species that produce both lactic acid and ethanol during primary fermentation. It is unclear how widespread this heterolactic fermentation phenotype is among ethanol-tolerant yeasts, but its presence in the fission yeast *S. japonicus* and budding yeasts like *H. vineae*, which are separated by ~1 billion years of evolution (22), may indicate that heterolactic fermentation is an ancient and conserved metabolic process among single-celled fungi. Arguing against this hypothesis is the lack of detectable lactic acid production by related strains of the same species, *e.g*., differences among *L. thermotolerans* isolates ((14) and data not shown). Whole genome sequencing and/or transcriptomic analysis of lactic acid-producing yeast during fermentation is needed to determine how the heterolactic fermentation occurs. Regardless, it is our hope that the strains described above and the primary souring process put forth here will add strength to the already growing sour beer movement in the U.S. and abroad.

## Acknowledgements

We thank Andrea Baillo, Elise Bochman, Henri Bochman, Chris Boggess, Kris Brown, Shannon Brown, Adam Covey, Sara Davidson, Jeff Ewer, Todd Green, Ted Herrera, Austin Kelley, Steve Llewelyn, Andrew Mason, Colin McCloy, Jared Miller, Ted Miller, Sasha Pefferman, Adam Quirk, Kevin Smolar, Jack Sramek, Caleb Staton, Julia van Kessel, and Linda van Kessel for collecting samples for our yeast hunting projects.

